# Comparing data-driven physiological denoising approaches for resting-state fMRI: Implications for the study of aging

**DOI:** 10.1101/2023.04.17.537223

**Authors:** Ali M Golestani, J. Jean Chen

## Abstract

Physiological nuisance contributions by cardiac and respiratory signals has a significant impact on resting-state fMRI data quality. As these physiological signals are often not recorded, data-driven denoising methods are commonly used to estimate and remove physiological noise from fMRI data. To investigate the efficacy of these denoising methods, one of the first steps is to accurately capture the cardiac and respiratory signals, which requires acquiring fMRI data with high temporal resolution. In this study, we used such high-temporal resolution fMRI data to evaluate the effectiveness of several data-driven denoising methods, including global-signal regression (GSR), white matter and cerebrospinal fluid regression (WM-CSF), anatomical (aCompCor) and temporal CompCor (tCompCor), ICA-AROMA. Our analysis focused on each method’s ability to remove cardiac and respiratory signal power, as well as its ability to preserve low-frequency signals and age-related functional connectivity (fcMRI) differences. Our findings revealed that ICA-AROMA and GSR consistently remove more heart-beat and respiratory frequencies, but also the most low-frequency signals. Our results confirm that the ICA-AROMA and GSR removed the most physiological noise at the expense of meaningful age-related fcMRI differences. On the other hand, aCompCor and tCompCor seem to provide a good balance between removing physiological signals and preserving fcMRI information. Lastly, methods differ in performance on young- and older-adult data sets. While this study cautions direct comparisons of fcMRI results based on different denoising methods in the study of aging, it also informs the choice of denoising method for broader fcMRI applications.

## Introduction

It has become established that for a method aiming to quantify brain function, resting-state blood-oxygenation level-dependent (BOLD) fMRI (rs-fMRI) metrics is exquisitely sensitive to underlying brain physiology, embodied in such variables as cerebrovascular reactivity (Chu et al., 2018; Golestani et al., 2016; Tsvetanov et al., 2020). Moreover, physiological contributions from cardiac and respiratory frequencies constitute a major source of noise in the blood-oxygenation level dependent (BOLD) fMRI data (Liu, 2016). Providing that the sampling rate of the fMRI data is sufficiently high to faithfully capture the fundamental cardiac and respiratory frequencies, the time-locked components of these noise sources can be largely reduced by a notch filter (Chen et al., 2019). However, the sampling rate of a typical fMRI data is not sufficiently high, resulting in aliasing of the physiological signals into lower frequencies, rendering the application of notch filters impractical. Alternatively, methods have been developed to model and remove phase-locked physiological on the BOLD fMRI signal based on recorded physiological signals (Glover et al., 2000). For these purposes, cardiac and respiratory recordings should be obtained during the fMRI data acquisition. However, this can be challenging, unreliable, and in some cases impossible due to experimental limitations (Agrawal et al., 2020). Data-driven methods represent an alternative, whereby estimates of the physiological effects are generated from the fMRI data itself.

One of the oldest data-driven denoising approaches is to regress out the global BOLD signal (GS), calculated by averaging signals of the voxels within the brain. Global-signal regression (GSR) assumes that because the effect of physiological signals on the fMRI data is widespread, global signal regression will remove physiological and other noise sources. However, GSR is controversial as it can also remove information relevant to brain function and connectivity (Liu et al., 2017). Brain states such as arousal and vigilance can also alter the GS through whole-brain effects (Gu et al., 2019; Wong et al., 2013). Nonetheless, GSR remains widely used for datasets with a high level of global noise (Chen et al., 2012), as it improves anatomical specificity of the connectivity maps (Fox et al., 2009) and increase the behavioral correlations with connectivity patterns (Li et al., 2019).

An alternative to GSR is to regress out the average signals derived from only the white matter (WM) and cerebrospinal fluid (CSF), where neuronal contributions are thought to be negligible (Bartoň et al., 2019). The WM-CSF regression approach found considerable application (He et al., 2020; Parkes et al., 2018; Satterthwaite et al., 2013; Scheel et al., n.d.). However, there is evidence that WM also contains information about brain function, especially at higher magnetic fields (Mazerolle et al., 2013). Moreover, the average signal across WM and CSF cannot account for regional-specific temporal variations of the physiological effects (Attarpour et al., 2021).

To address this issue, the CompCor family of methods applies principal component analysis (PCA) on a collection of signals from these “noise” regions of interest (ROIs) to decompose them into uncorrelated components, such that only a specific number of components with the highest variance are removed (Behzadi et al., 2007). CompCor has two variants; anatomical CompCor (aComCor) defines noise sources anatomically, by focusing on signals within WM and CSF anatomical masks, whereas temporal CompCor (tCompCor) defines noise sources temporally, by focusing on signals with high temporal standard deviation irrespective of their spatial origin. The aCompCor method is built into such software packages as the CONN Toolbox (Whitfield-Gabrieli and Nieto-Castanon, 2012), and has been widely applied.

As a departure from these conventional methods, independent component analysis (ICA) has been used to spawn a family of techniques for extracting representatives of the physiological noise from the fMRI data (Glasser et al., 2018; Golestani and Chen, 2022; Thomas et al., 2002). ICA decomposes the fMRI data into spatially independent components, and assuming that the physiological noise and neuronally driven signals are spatially independent, ICA can separate them into different components. The noise-related components can be manually identified based on their spatial, temporal, and spectral features, especially as physiological noise can manifest as semi-regular head motion with distinct spatial patterns. Nonetheless, this process is subjective, which can introduce inter-cohort and inter-study variability. To address this issue, a series of ICA variants, such as ICA-FIX (semi-automatic noise classification) and ICA-AROMA (Automatic Removal of Motion Artifacts) have been developed that allow noise-related components to be spatially and temporally identified *a priori (Pruim et al., 2015b; Salimi-Khorshidi et al., 2014)*. Thus, an advantage of ICA-AROMA over ICA-FIX is that the former makes use of spatiotemporal features to identify noise components, and thus does not require training of the noise classifiers with each new data set while retaining much of the functionally relevant correlational structure structure in the data. The performance of ICA-AROMA has been compared favorably against that of ICA-FIX (Dipasquale et al., 2017; Pruim et al., 2015a), and ICA-AROMA has become increasingly adopted in rs-fMRI analysis (Cohen et al., 2021; Dipasquale et al., 2017).

Despite widespread application of the data-driven noise removal methods, the efficacy of these noise removal techniques is unclear. A systematic evaluation of efficacy faces a number of challenges. First, since the typical sampling rate of the fMRI data is above the Nyquist frequency of the respiration and heartbeat, these physiological signals alias into low frequencies, and therefore investigating the efficacy of the noise removal techniques is challenging. Thus, it is typically impossible to quantify the amount of major physiological noise sources in the signal. Second, the biggest roadblock for the application for rs-fMRI is in clinical translation, and it remains unclear whether superior sensitivity or repeatability translate into superior reflection of biological differences. Third, when assessing biological differences using rs-fMRI functional connectivity (fcMRI), there is no ground truth to compare against. In this regard, the sole dependence on conventional MRI quality metrics, such as sensitivity and reproducibility, may not be ideal for rs-fMRI, as the latter is expected to be variable with time (Aedo-Jury et al., 2020; Tailby et al., 2015).

In this study, to address these challenges, we adopt the following methodological choices. First, we devised a pseudo-ground truth for assessing physiological noise content, namely, by examining the BOLD signal’s frequency spectra. This approach is facilitated by our high temporal-resolution fMRI data and simultaneously recorded physiological signals to accurately assess the location and contribution of physiological frequencies. Second, physiological contributions may be linked to head motion and may vary with age (Makedonov et al., 2013; Tsvetanov et al., 2015; Van Dijk et al., 2012), as it was previously found to vary with Alzheimer’s disease (Li et al., 2021). Thus, by introducing the variable of age into the study, we can observe the impact of age on denoising performance. Furthermore, instead of relying on sensitivity and reproducibility as metrics of denoising quality, we compared denoising methods by the extent to which the output of each denoising technique can preserve age-related functional-connectivity (fcMRI) differences. We use these approaches to evaluate all data-driven denoising methods available through the commonly used fMRIPrep pipeline. We hypothesized that methods that removed more low-frequency signal power also resulted in a loss of sensitivity to age-related resting-state fcMRI (rs-fcMRI) differences.

## Method

### Participants and data acquisition

18 healthy young subjects (age=26.7 ± 6.5 years) and 18 healthy older subjects (age=74.2 ± 7.0 years) were imaged using a Siemens TIM Trio 3T scanner (Siemens Healthineers, Erlangen, Germany). All participants provided written informed consent as per the policy of our institutional research ethics board. rs-fMRI scans were collected using simultaneous multi-slice (SMS) echo-planar imaging (EPI) BOLD (TR/TE = 380/30 ms, FA = 40°, 20 5-mm slices, 64×64 matrix, 4×4×5 mm voxels, multiband factor = 3, 1,950 volumes). During each scan, cardiac pulsation was recorded using the scanner pulse oximeter, whereas the respiratory signal was recorded using a Biopac^TM^ system (Biopac Systems Inc. California, USA). A T1-weighted 3D anatomical data set (1mm isotropic resolution) was also acquired for each participant.

### Data preprocessing and physiological denoising

The anatomical segmentation was performed using fMRIPrep, as shown in Fig. 1. Specifically, the T1 anatomical images were skull-stripped using a Nipype implementation of Advanced Normalization Tools (ANTs) (Avants et al., 2011), following which tissue segmentation was performed using FMRIB Software Library (FSL) and spatial normalization to standard MNI152 space was performed using ANTs. This step resulted in grey matter (GM), WM and CSF segmentations for each data set.

fMRI data preprocessing was performed using fMRIPrep (Esteban et al., 2019), and includes motion correction, spatial smoothing (5mm FWHM), high-pass filtering (>0.01 Hz) and brain extraction. This is a common processing pipeline applied prior to all physiological denoising methods (no correction). The preprocessed data was channeled separately through five data-driven denoising methods for comparison, as implemented through fMRIPrep. These methods include:

- Global signal regression (GSR): Global signal is generated by averaging signals within the brain mask (excluding CSF).
- White matter and CSF signal regression (WM-CSF): WM and CSF ROIs were obtained from the segmentation of the T1-weighted anatomical image of each subject, as explained above. WM and CSF signal is generated by averaging the signals within the anatomically-derived eroded masks.
- Anatomical CompCor (aCompCor): The five principal components (PCs) with the highest explained variance are used as confounders.
- Temporal CompCor (tCompCor): All PCs identified by tCompCor are regressed out.
- AROMA (ICA-AROMA): Based on ICA, AROMA exploits a small set of four robust theoretically motivated temporal and spatial features associable to head motion, and all independent components exhibiting these features are identified as nuisance and removed. Though devised for head-motion correction, AROMA has found success for physiological denoising more broadly (Griffanti et al., 2017)
- No correction: All methods were also compared to the case of no physiological denoising for reference.

Regressors of each method are estimated using fMRIprep (Esteban et al., 2019), publicly available at fmriprep.org.

**Figure 1.**
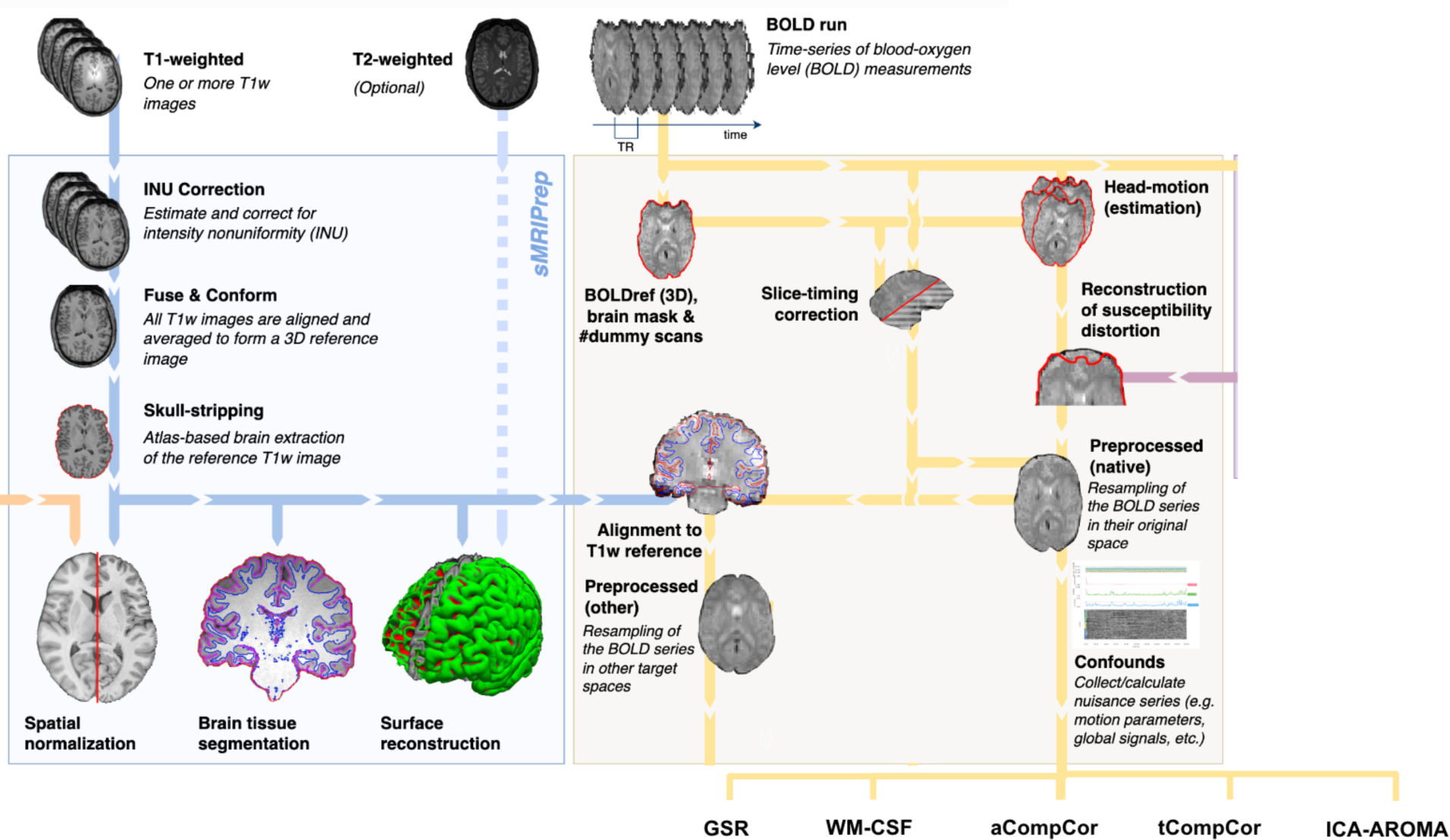
Summary of preprocessing steps. (figure adapted from www.fmriprep.org). All illustrated preprocessing steps were applied to the fMRI and T1-weighted data as relevant. The tissue segmentation required for denoising is performed through FreeSurfer. In particular, the fMRI data underwent slice-timing correction, alignment to the T1 images, head motion estimation, susceptibility-distortion correction and confound estimation. The confounds produced by this pipeline are in turn used by some of the denoising strategies.

**Figure 1.**
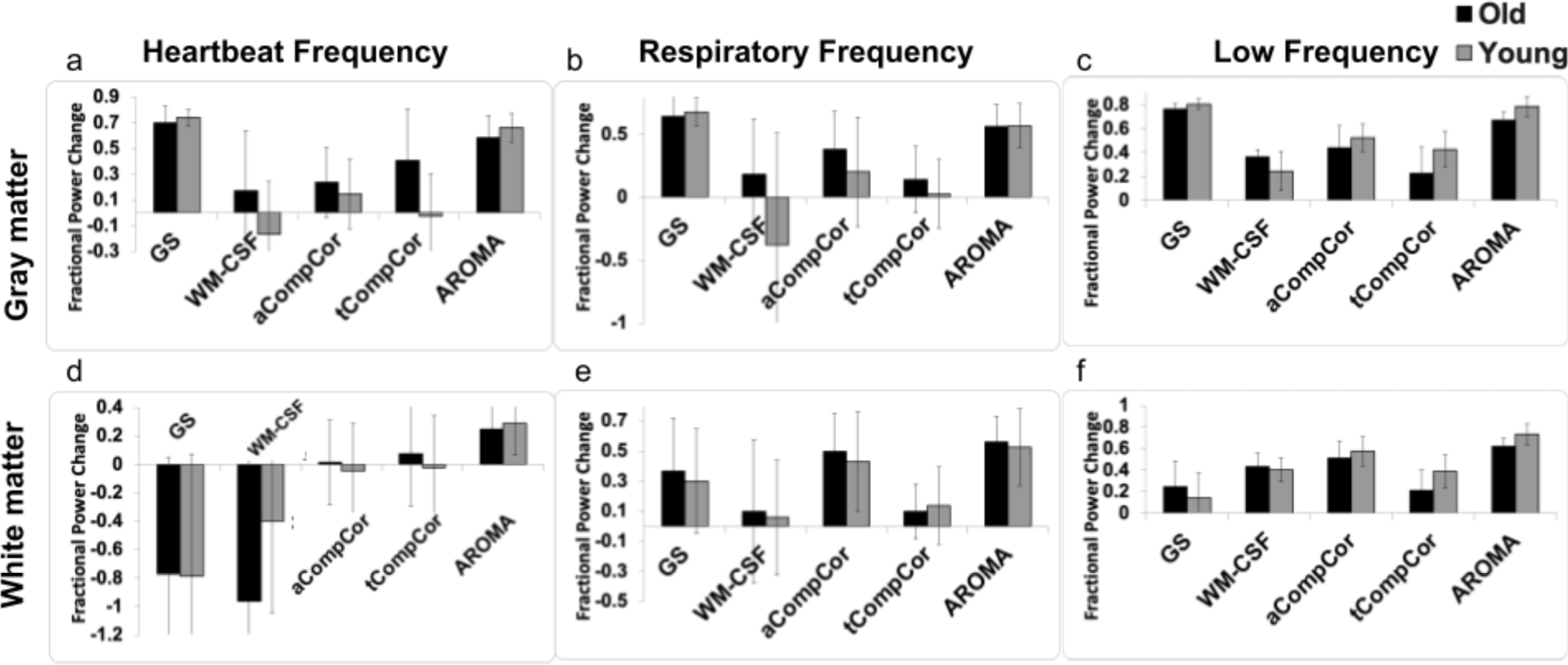
The effect of denoising strategies on the BOLD signal spectral power in young and older adults. Fractional power changes in cardiac (a & d), respiratory (b & e), and low-frequency bands (c & f) for the gray matter (a, b, c) and white matter (d, e, f) signals. Young and older adults are shown gray and black, respectively. Note that a positive power change represents a reduction in the spectral power due to denoising, and vice versa. Error bars represent the group-wise standard deviation.

### Evaluation metrics

#### BOLD-signal spectral power

In order to identify the BOLD signal spectral peaks associated with time-locked physiological processes, subject-specific heartbeat and respiration frequencies were estimated based on the peak in the spectrum of their physiological recordings. Then, for each data set, the fMRI signal is averaged separately across the gray matter and white matter, respectively, for each individual and the spectrum of the averaged signals is calculated. We computed the total fMRI signal power pre- and post-denoising in three frequency bands:

1. Cardiac band: a 0.1 Hz-wide band centered at the subject-specific heartbeat;
2. Respiratory band: a 0.2 Hz-wide band centered around each subject’s respiration frequency;
3. Low-frequency band: the frequency band between 0.01 and 0.1 Hz, commonly used for fcMRI assessments.

A successful denoising technique should maximally remove the frequencies associated with the cardiac and respiratory bands while preserving information in the low-frequency band. To evaluate the extent to which each denoising method alters the power contribution of these frequency bands, we computed the fractional power change as

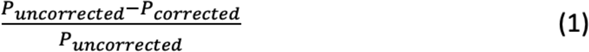

where *P*_uncorrected_ is the power of the fMRI signal before noise correction in one of the three frequency bands and *P*_corrected_ is the power of the fMRI signal after noise correction in these frequency bands. The use of fractional spectral-power change allows direct comparison across methods and age groups.

#### rs-fcMRI metrics

For each processed fMRI dataset, template-based rotation (TBR) was used to generate fcMRI maps using Yeo functional network parcellations templates (Yeo et al., 2011)). TBR is a new analytic technique that was designed for utilizing a priori functional parcellations to guide the analysis of individual sessions. TBR is similar to dual regression, but reverses the direction of prediction such that instead of individual volumes (time points) being predicted as linear sums of templates, templates are predicted as linear sums of volumes (Schultz et al., 2014). The TBR step produced fcMRI strengths quantified as t-maps. Subsequently, for each network, a voxel-wise 2-factor repeated-measures analysis of variance (ANOVA) is used with processing method and age as the factors. The F-map associated with the factor of age is considered as a pseudo-ground truth, as it reflects fcMRI differences between age groups regardless of the denoising method. Next, for each network and denoising method, an age-related fcMRI difference map is generated using voxel-wise unpaired t-tests, and significance is determined at the cluster-significance threshold (Woo et al., 2014). The performances of the denoising methods in terms of fcMRI maps are lastly evaluated by comparing the difference t-map created by each method with the pseudo-ground truth F-map.

These fcMRI age differences relative to the pseudo-ground-truth age difference are further evaluated quantitatively through three metrics. First, for each of the 7 networks, the correlations between the pseudo-ground-truth F-maps and the absolute value of the per-method t-map were calculated. Secondly, to account for the fact that we may not be able to assume similar distributions of the t- and F-statistics, cosine similarity between the t- and F-maps were also computed to assess similarity. Thirdly, spatial overlap was computed using the Dice coefficient. Specifically, F-maps and t-maps are both thresholded at p = 0.01, uncorrected for multiple comparisons. Third, significant cluster sizes at the p <0.05 level were estimated using FSL *cluster*, and the corresponding cluster sizes were applied to the thresholded F- and t-maps to achieve a significant cluster threshold equivalent of p = 0.05. Dice coefficients between the thresholded F-map and the t-map of each method are calculated.

All performance metrics, including the fractional power change, the correlation coefficients, the cosine-similarity and Dice indices, are then compared between pairs of methods using the Wilcoxon signed-rank test.

## Results

In Fig. 1 are shown the fractional power changes in the cardiac-frequency, respiration-frequency and low-frequency bands for the whole GM (Fig 1a, b, c) and WM signals (Fig. 1d, e, f). Results for young and older adults are shown in gray and black, respectively. In the GM, all denoising methods resulted in spectral-power reduction in all three frequency bands of the BOLD signal, removing as much as 60-70% of the signal power in the respiratory and cardiac bands. GSR and AROMA removed the highest percentages of cardiac and respiratory BOLD-signal power, followed by aCompCor. However, GSR and AROMA also removed the largest percentages of the BOLD signal from the low-frequency band (up to 80%). In the WM, all methods appear less effective in removing cardiac power, with AROMA removing the most signal from (>70%). These differences are statistically significant, as shown in Table 1.

**Table 1.**
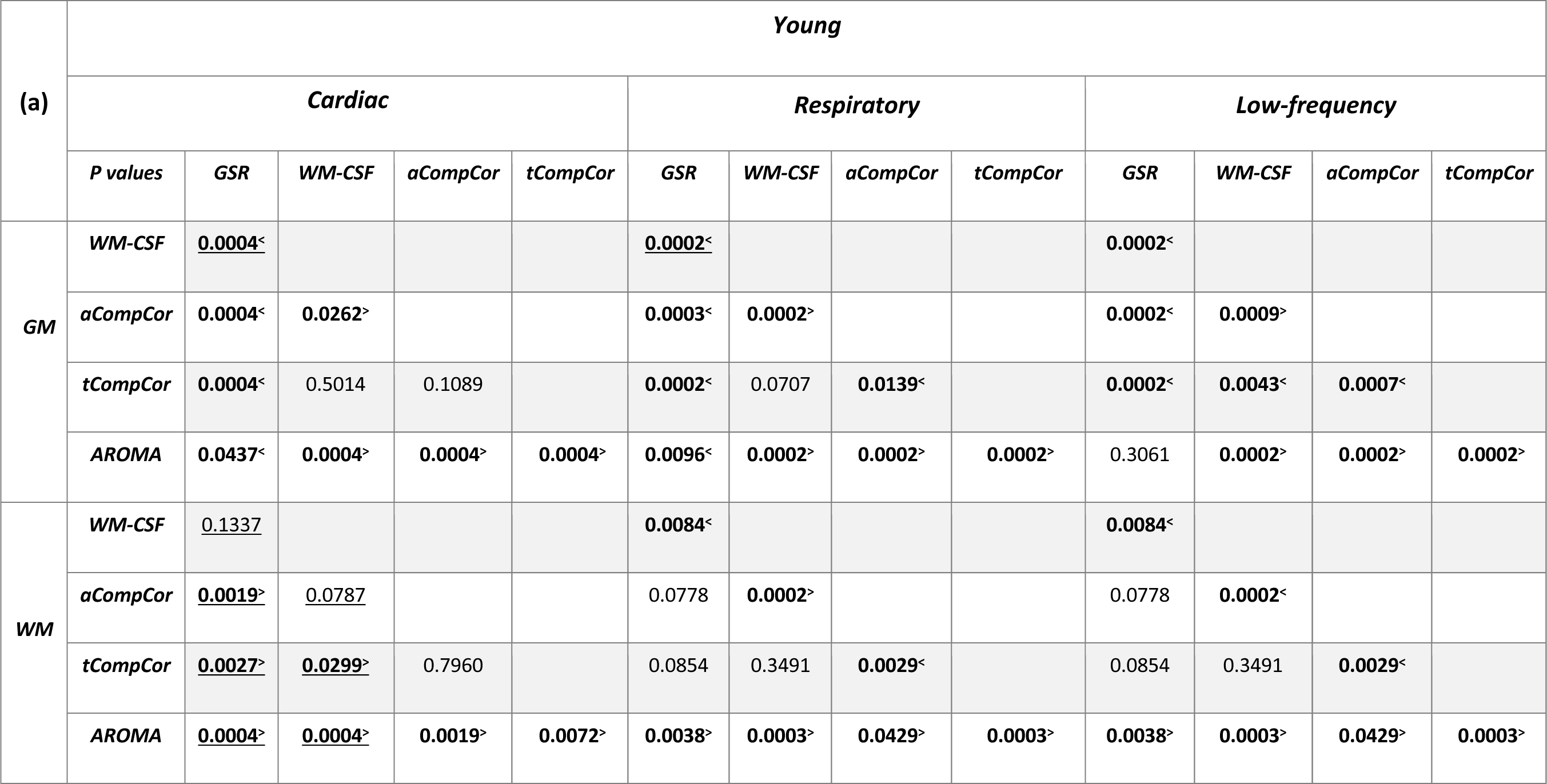
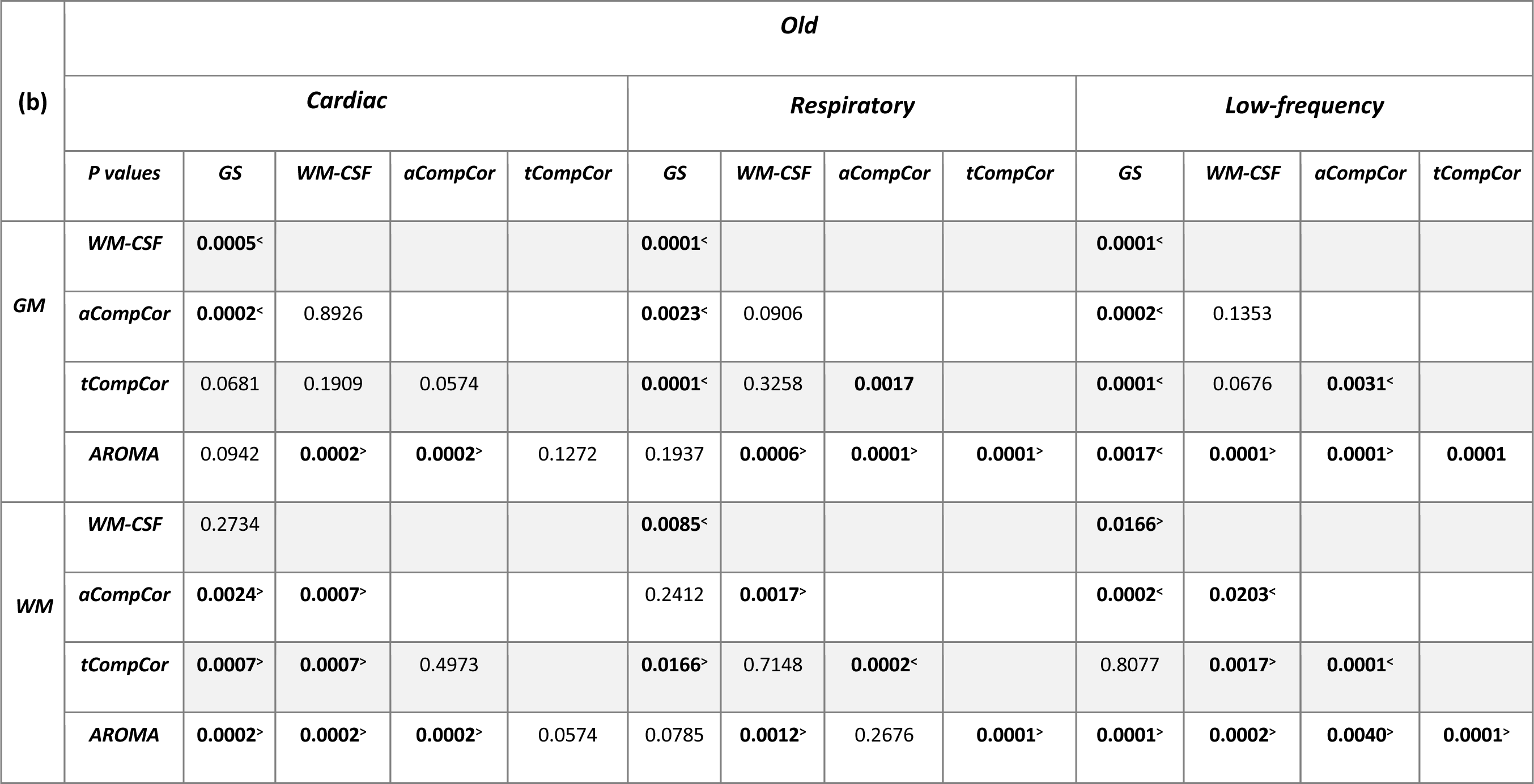
Pairwise comparison of fractional power removed associated with different denoising methods. (a) Data from young cohort; (b) data from older cohort. P values for GM and WM pairwise comparisons in different frequency bands, with those that are statistically significant shown in bold face. Underlining indicates where GSR and WM-CSF resulted in an increase rather than a decrease in spectral power. ‘<’ indicates where the denoising method on the row label removed a lower fractional power than the method on the column label, and ‘>’ indicates the opposite.

Taking Fig. 1 and Table 1 in conjunction, we observe that In young adults, for GM, GSR and AROMA removed significantly more cardiac noise than all methods, and more respiratory noise than all but WM-CSF regression. GSR and AROMA removed more low-frequency signal than all other denoising methods. For WM, AROMA removed the most cardiac, respiratory and low-frequency power, followed by aCompCor. GSR removed the least low-frequency power in the WM. GSR and WM-CSF regression introduced increases to cardiac power. For the older-adult data, WM-CSF regression performed worse, adding to rather than removing power from the cardiac and respiratory bands. Other trends are similar as in the young group. Moreover, across all noise bands, all methods appear to remove less noise power from the BOLD signals of young adults compared to those of older adults. Conversely, all methods removed more low-frequency power from the data of young adults. Nevertheless, the difference between young adults and older adults was not statistically significant. Across both age groups, aCompCor and tCompCor performed similarly for removing cardiac noise, but aCompCor is more effective than tCompCor for removing respiratory noise.

Fig. 2 summarizes the fcMRI values (TBR-based correlation t statistic) corresponding to the data in Fig. 1. The network-specific fcMRI measurements are represented as box plots generated from the 7 network templates defined by the Yeo fcMRI atlas. Thus, these resultant values do not account for differences in the network spatial extents resulting from the different denoising methods. The GSR and AROMA methods resulted in the greatest reductions in fcMRI values, in all but the limbic network. The GSR method also visibly reduced the age difference in fcMRI, particularly in the visual network.

**Figure 2.**
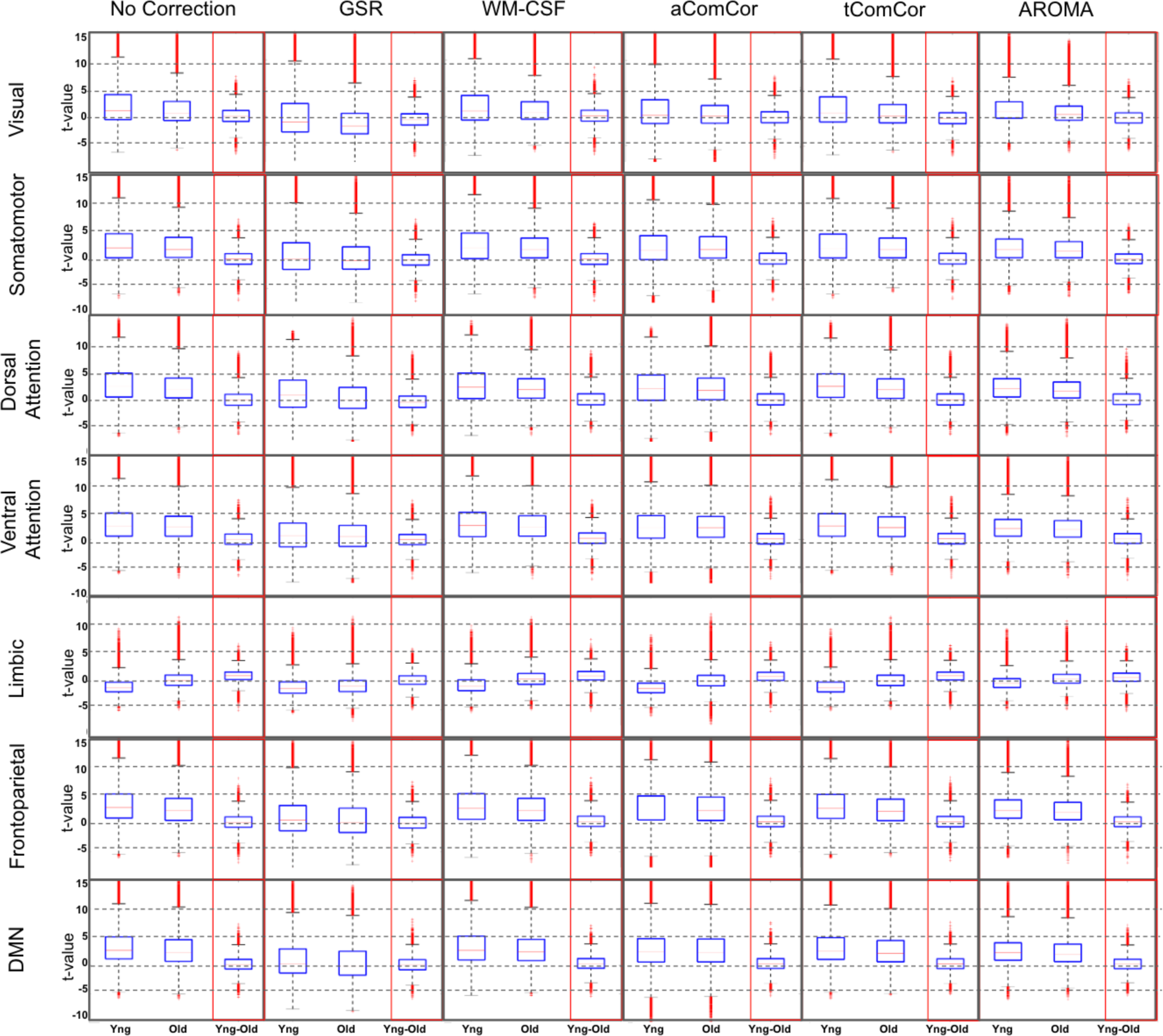
Comparison of fcMRI measurements corresponding to different denoising methods and different networks. The network-specific fcMRI measurements are represented as box plots generated from the 7 network templates defined by the Yeo fcMRI atlas, labeled along the vertical axis. The mean fcMRI values are plotted for the young (Yng), older (Old) and age difference (Yng-Old). DMN = default-mode network.

In Fig. 3 we demonstrate the age-related fcMRI differences generated for the case of the default-mode network (DMN as an example) using different denoising methods. The top row shows the F-maps associated with significant age-related differences, regardless of denoising method, considered as the pseudo ground-truth. The age-related fcMRI difference t-maps generated using different denoising methods are also shown in lower rows of Fig. 3. Qualitatively, the maps generated from aCompCor, tCompCor and WM-CSF appear to be the most similar to the pseudo ground truth map.

**Figure 3.**
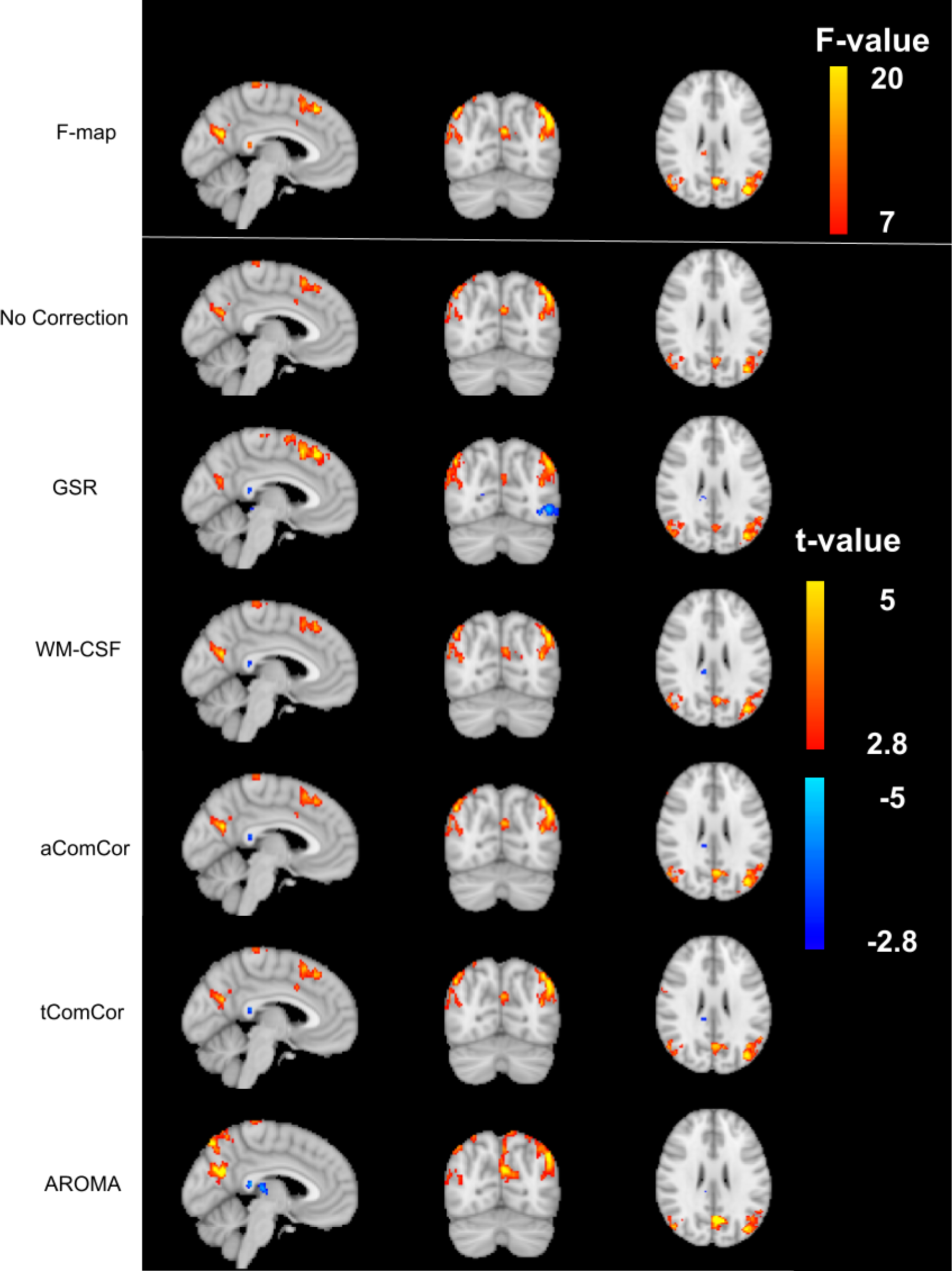
Sample age-related connectivity-difference maps for the default-mode network. The F-map corresponding to the age effect is shown on top to represent the pseudo ground-truth age effect on fcMRI strength. Each row shows the difference t-map generated from each of the denoising methods. The color bar represents the range of F-values for the F-map and t-values for the t-maps.

Fig. 4 shows the spatial correlation values, cosine similarity and Dice coefficients calculated between the pseudo ground-truth F-map and the t-map for different processing methods, averaged across the 7 resting-state networks. The statistical significance of pairwise method comparisons associated with the values presented in the figures are shown in Table 2, respectively. Based on these metrics, the age-related fcMRI-difference maps generated using AROMA and GSR denoising are the least similar to the ground-truth maps (lowest p values). In contrast, the maps associated with the aCompCor and tCompCor methods exhibit the highest similarity with the pseudo-ground-truth maps (highest p values).

**Figure 4.**
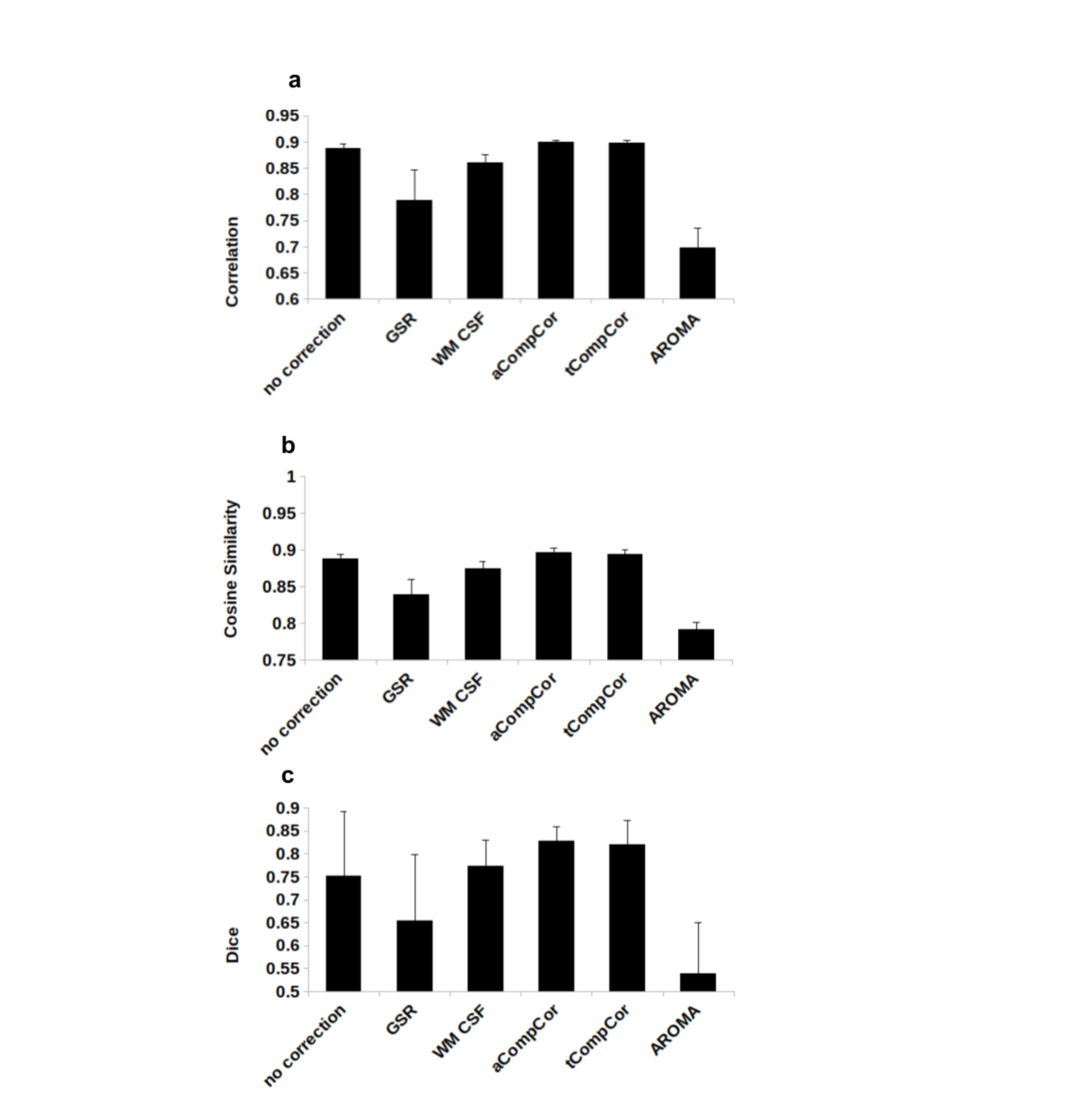
Quantitative comparisons of age-related fcMRI differences obtained through all denoising methods across 7 resting-state networks. (a) The spatial correlation between age-related differences are assessed by correlating the pseudo-ground-truth F-maps and the per-method t-maps, averaged across all resting-state networks. Error bars represent the group-wise standard deviation. Compared to the maps with no correction, those generated with aCompCor and tCompCor show an increase in the similarity to the pseudo-ground-truth map, whereas those generated with AROMA and GSR show a reduced similarity. A similar pattern is found through the cosine similarity metric (b). (c) The Dice coefficient computed between the thresholded pseudo-ground-truth F-map and the difference t-map between age groups, averaged across the 7 resting-state networks. Compared to the maps with no correction, those generated with AROMA and GSR show reduced similarity with the pseudo-ground-truth maps.

**Table 2.** Statistical significance of inter-method comparisons of the agreement in fcMRI spatial extent between the method-specific result and the pseudo-ground truth. The p-values displayed correspond to the correlation values in Fig. 3a, and were generated using the Wilcoxon signed-rank test comparing spatial correlation between age-related differences through the pseudo-ground-truth F-map and the per-method t-maps.

|  | *Correlation Coefficient* |  |  |  |  | *Cosine similarity* |  |  |  |  | *Dice coefficient* |  |  |  |  |
| --- | --- | --- | --- | --- | --- | --- | --- | --- | --- | --- | --- | --- | --- | --- | --- |
| *p values* | *No Corr.* | *GS* | *WM-CSF* | *aCompCor* | *tCompCor* | *No Corr.* | *GS* | *WM-CSF* | *aCompCor* | *tCompCor* | *No Corr.* | *GS* | *WM-CSF* | *aCompCor* | *tCompCor* |
| **GS** | 0.016 |  |  |  |  | 0.016 |  |  |  |  | 0.016 |  |  |  |  |
| **WM-CSF** | 0.016 | 0.016 |  |  |  | 0.032 | 0.016 |  |  |  | 0.578 | 0.016 |  |  |  |
| **aCompCor** | 0.016 | 0.056 | 0.0156 |  |  | 0.016 | 0.016 | 0.016 |  |  | 0.016 | 0.016 | 0.016 |  |  |
| **tCompCor** | 0.016 | 0.056 | 0.0156 | 0.4688 |  | 0.016 | 0.016 | 0.016 | 0.375 |  | 0.016 | 0.016 | 0.016 | 0.578 |  |
| **AROMA** | 0.016 | 0.056 | 0.0156 | 0.0156 | 0.0156 | 0.016 | 0.016 | 0.016 | 0.016 | 0.016 | 0.016 | 0.031 | 0.016 | 0.016 | 0.016 |

## Discussion

In this work, we assessed the influence of a variety of popular data-driven denoising methods on resting-state BOLD signal-power and on rs-fcMRI maps in commonly observed resting-state functional networks. We further assess the impact of denoising choice on the resultant age-related fcMRI differences in these networks. Using a fMRI dataset acquired at high sampling rate, we were able to directly calculate the amount of spectral power altered by each method at each subject’s cardiac and respiratory frequencies. To summarize, in the GM, all methods reduced the BOLD spectral power in these physiological frequency ranges, albeit the degree of removed spectral power is significantly different among the methods, with GS and AROMA removing the most power. These methods, however, also removed the greatest amount of low-frequency power, which, amongst the three frequency bands, pertains the most to resting-state brain connectivity. In contrast, CompCor (aCompCor and tCompCor) seems to offer a good compromise between removing physiological noise and retaining fcMRI-related information. In the WM, all methods reduced the signal power in the respiratory frequency ranges. However, the performance of the methods in removing cardiac frequencies is more variable across subjects (see Fig. 1). While our findings address age differences, they can also be applied to fcMRI studies in disease conditions.

### Physiological noise power spectrum

Cardiac pulsation and respiration are the two chief sources of physiological noise in the fMRI signal, especially problematic in resting-state fMRI. Cardiac pulsations generate localized brain motion as well as creating inflow effects in and around blood vessels and CSF. On the other hand, thoracic movement during breathing results in magnetic field alteration and phase shift in the image. Moreover, respiration induces fluctuation in the level of arterial CO_2_, which has a vasodilating effect and can induce changes in the fMRI BOLD signal. The frequency of physiological signals in an adult human typically falls within the range of 0.2 to 0.3 Hz for respiratory and 0.8 to 1.2 Hz for cardiac signals. In a conventional fMRI scan with a TR of 2 seconds, these signals alias into the low-frequency range (0.01 to 0.1 Hz), which is known to contain information about fcMRI (Cordes et al., 2001). Physiological denoising in a conventional fMRI dataset may result in a decrease in signal power in the low-frequency range. However, it is impossible to determine whether the removed power is related to physiological fluctuations or neuronal connectivity.

### Choice of denoising methods

For context, the methods evaluated in this study are among the most commonly used in the resting-state fMRI literature. Specific to the study of the effect of aging on resting-state connectivity, all of the denoising methods have found broad application, including GSR (Betzel et al., 2014; Chan et al., 2014; Geerligs et al., 2015; Siman-Tov et al., 2016; Stumme et al., 2020; Zhang et al., 2014), WM-CSF (Betzel et al., 2014; Chan et al., 2014; Farras-Permanyer et al., 2019; Grady et al., 2016; Koch et al., 2010; Mancho-Fora et al., 2020; Siman-Tov et al., 2016; Song et al., 2014; Varangis et al., 2019; Xie et al., 2020; Zhang et al., 2014; Zhong and Chen, 2022), CompCor (Hamada et al., 2021; Hausman et al., 2020; Onoda and Yamaguchi, 2013; Patil et al., 2021; Podgórski et al., 2021), and ICA-AROMA (Stumme et al., 2020). Moreover, some recent studies of aging used no physiological denoising (Ai et al., 2020; Damoiseaux et al., 2008; Huang et al., 2015).

### Impact on BOLD signal spectrum

Owing to the high sampling rate of our fMRI data, we were able to dissect the differential effect of different denoising methods by examining the resultant fcMRI signal power spectrum. All of the investigated methods are shown to reduce the spectral power in the cardiac and respiratory frequencies of the GM signals. All methods resulted in spectral-power reduction in all three frequency bands of the gray matter signal, removing as high as 60-70% of the signal in the respiratory and cardiac bands. The amount of the removed power spectrum is vastly different among the methods, with GS and AROMA removing the most power followed by aCompCor. Taking GM and WM together, AROMA also removes more signal power in the physiological frequency bands compared to all other denoising methods. AROMA, developed for identified head motion, appears to capture physiologically driven head motion quite well, constituting a promising ICA-based method of physiological denoising, particularly one without the need for training and manual intervention.

Compared to the GM, the results in the WM are more variable across subjects. Notably, among all the methods, only AROMA appeared to reduce cardiac power (Fig. 1d). All methods reduced power in the respiratory and low-frequency frequency bands (Fig. 1e & f), with aCompCor and AROMA leading the other methods in this regard. Nevertheless, the power-reduction variability in WM is much larger than that in GM, especially for GSR, WM-CSF and aCompCor. GSR is also very effective at suppressing the contribution of physiological frequencies in the GM, but not very effective in the WM. Unsurprisingly, both GSR and WM-CSF regression introduced increases to cardiac power, and are thus inappropriate for WM fMRI denoising. This observation could be attributed to the manner in which the noise regressors are derived for these methods, in which the WM signal is indiscriminately designated as noise. Interestingly, WM-CSF did not significantly reduce WM BOLD signal power in the cardiac and respiratory power, although this method can be viewed as involving regressing the WM signal out of itself. Although aCompCor also derives regressors from WM, its performance is superior to that of WM-CSF, as it only designated the leading principal components of the WM signal (and CSF) signal as noise regressors (Fig. 1e).

### Impact on fcMRI strength

While GS and AROMA perform better in removing the physiological noise power spectrum, they also remove the most BOLD signal power from the low-frequency band, as they do not discriminate “signal” from “noise” through location or signal variance. This specifically concerns the low-frequency band, the established location for the majority of the neurovascular contribution to brain functional connectivity (D Cordes 1, V M Haughton, K Arfanakis, J D Carew, P A Turski, C H Moritz, M A Quigley, M E Meyerand, 2001). However, low-frequency variability in the respiratory and cardiac pattern can also induce BOLD signal variations in this frequency band (Birn et al., 2006; Chang et al., 2013; Wise et al., 2004). Thus, based on the power spectrum alone, it is impossible to understand if the removed low-frequency spectrum is related to the brain connectivity or physiological noise. To address this, we calculated rs-fcMRI maps and compared the methods based on their abilities in representing age-related rs-fcMRI differences. Our results confirmed that the removed low-frequency signals by GS and AROMA do in-fact carry contribution from meaningful age effects on fcMRI measures. Specifically, GSR indiscriminately removes the global-mean signal, which, though driven strongly by respiration (Power et al., 2017), obviously also contains neuronally relevant signal, as GSR is also seen to reduce the ability to fully capture fcMRI amplitude and age-related effects compared to other methods (Fig. 2). Nonetheless, some of the respiratory signals may still be entangled with both head motion and neuronal activity (Shams et al., 2021; Yuan et al., 2013). In this regard, limiting noise regressors to originate from the WM and CSF appears a more cautious approach. By the same measures, aCompCor and tCompCor generate maps with the highest similarity to the ground truth, as demonstrated by spatial correlation, cosine similarity, and Dice coefficient (Fig. 4 **and** Tables 2).

### Role of age

The performances of different denoising methods vary by age. All methods removed less noise power in the young adults. This observation can be explained by the method we calculated the relative power removed. The amplitude of low-frequency fluctuation decreases with advancing age (Hu et al., 2014); one possible reason is that age-related vascular stiffness may reduce the contribution of vascular modulation to the fMRI signal. Thus, one can expect the low-frequency band to contribute less to the total rs-fMRI signal power, and the denominator of Eq. (1) to be lower in the older adults, resulting in higher relative noise power change by denoising. Age-related decrease of the low-frequency power can attribute to decline in brain function during aging.

In aging studies of fcMRI, rs-fcMRI reduction in the DMN is the most commonly reported finding (for a review please refer to (Jockwitz and Caspers, 2021)), although the extent and location of the connectivity reduction differ among the studies, consistent with our finding (Fig. 2). For instance, Stumme et al. (Stumme et al., 2020) used AROMA denoising and reported reduced connectivity in the visual network and no changes in the DMN and somatomotor networks, whereas other studies that used tCompCor or WM-CSF denoising have reported no change (Zhang et al., 2014) or reduced (Chan et al., 2014; Onoda and Yamaguchi, 2013; Zhang et al., 2014) connectivity in the visual network and reduced connectivity in the DMN (Zhang et al., 2014)(Betzel et al., 2014; Chan et al., 2014; Grady et al., 2016; Onoda and Yamaguchi, 2013), as well as no connectivity change in the somatomotor network (Betzel et al., 2014; Chan et al., 2014; Onoda and Yamaguchi, 2013). Here, we demonstrate that one possible source of the variation across studies is the choice of method for physiological denoising. Despite the fact that the spectral power removed by different denoising methods were at times not significantly different, these differences resulted in observable differences in the resultant age-related fcMRI differences. Specifically, as shown in Fig. 2, the GSR method also visibly reduced the age difference in fcMRI, particularly in the visual network, while CompCor and WM-CSF minimally impact the observable age difference in connectivity. This finding is consistent with CompCor methods revealing age differences that are the most consistent with the pseudo ground-truth.

### Limitations

This study presents several limitations. First, we focused only on the denoising methods implemented in fmriprep. More advanced methods have been developed recently (Agrawal et al., 2020; Bancelin et al., n.d.; Shin et al., 2022) we chose to compare only the methods implemented in fmriprep, as they may be more commonly used. The same methodology can be used to compare other physiological denoising techniques. Moreover, we only evaluated the methods in their ability to remove physiological signals.

The methods can perform differently in removing other types of noise. For example, ICA-AROMA has been shown to be more successful in removing head motion compared to aCompCor (Pruim et al., 2015a). In the absence of a ground-truth, we developed a pseudo-ground-truth by generating the age-difference maps from all denoising methods. The F-map shows brain regions with significantly different connectivity between the age groups, regardless of the denoising method. It is important to acknowledge that the pseudo-ground-truth can have limitations. The accuracy of the pseudo-ground-truth map can be influenced by the denoising methods used. If the denoising methods themselves are biased, this can affect the accuracy of the F-map. It may be useful to compare results across multiple denoising methods to assess the consistency of findings and identify potential sources of bias. Overall, it is important to interpret these results with caution.

## Conclusion

In this study, we compared the outcomes of several data-driven noise removal methods in removing power spectrum in the physiological noise while preserving age-related differences in functional connectivity. We observed that there is a trade-off between the two. Methods like GSR and AROMA excel in removing physiological noise, but at the cost of removing brain connectivity information. In comparison, aCompCor and tCompCor remove physiological noise effectively while also retaining neuronal information. The findings of this study can inform the choice of rs-fMRI denoising methods. It can also affect the interpretation of age effects on rs-fMRI metrics, suggesting that age effects on rs-fMRI signal power and fcMRI metrics calculated following different denoising methods cannot be directly compared.

## Acknowledgements

The authors would also like to acknowledge financial support from Canadian Institutes of Health Research (JJC). We also thank Mr. Jonathan Kwinta for assistance in data collection.

## References

1. Aedo-Jury, F., Schwalm, M., Hamzehpour, L., Stroh, A., 2020. Brain states govern the spatio-temporal dynamics of resting-state functional connectivity. Elife 9. https://doi.org/10.7554/eLife.53186

2. Agrawal, U., Brown, E.N., Lewis, L.D., 2020. Model-based physiological noise removal in fast fMRI. Neuroimage 205, 116231.

3. Attarpour, A., Ward, J., Chen, J.J., 2021. Vascular origins of low-frequency oscillations in the cerebrospinal fluid signal in resting-state fMRI: Interpretation using photoplethysmography. Hum. Brain Mapp. https://doi.org/10.1002/hbm.25392

4. Avants, B.B., Tustison, N.J., Song, G., Cook, P.A., Klein, A., Gee, J.C., 2011. A reproducible evaluation of ANTs similarity metric performance in brain image registration. Neuroimage 54, 2033–2044.

5. Bancelin, D., Bachrata, B., Bollmann, S., de Lima Cardoso, P., Szomolanyi, P., Trattnig, S., Robinson, S.D., n.d. Unsupervised physiological noise correction of fMRI data using phase and magnitude information (PREPAIR). https://doi.org/10.1101/2022.02.18.480884

6. Bartoň, M., Mareček, R., Krajčovičová, L., Slavíček, T., Kašpárek, T., Zemánková, P., Říha, P., Mikl, M., 2019. Evaluation of different cerebrospinal fluid and white matter fMRI filtering strategies— Quantifying noise removal and neural signal preservation. Human Brain Mapping. https://doi.org/10.1002/hbm.24433

7. Behzadi, Y., Restom, K., Liau, J., Liu, T.T., 2007. A component based noise correction method (CompCor) for BOLD and perfusion based fMRI. Neuroimage 37, 90–101.

8. Betzel, R.F., Byrge, L., He, Y., Goñi, J., Zuo, X.-N., Sporns, O., 2014. Changes in structural and functional connectivity among resting-state networks across the human lifespan. Neuroimage 102 Pt 2, 345– 357.

9. Birn, R.M., Diamond, J.B., Smith, M.A., Bandettini, P.A., 2006. Separating respiratory-variation-related fluctuations from neuronal-activity-related fluctuations in fMRI. Neuroimage 31, 1536–1548.

10. Chang, C., Metzger, C.D., Glover, G.H., Duyn, J.H., Heinze, H.-J., Walter, M., 2013. Association between heart rate variability and fluctuations in resting-state functional connectivity. Neuroimage 68, 93– 104.

11. Chan, M.Y., Park, D.C., Savalia, N.K., Petersen, S.E., Wig, G.S., 2014. Decreased segregation of brain systems across the healthy adult lifespan. Proc. Natl. Acad. Sci. U. S. A. 111, E4997–5006.

12. Chen, G., Chen, G., Xie, C., Ward, B.D., Li, W., Antuono, P., Li, S.-J., 2012. A method to determine the necessity for global signal regression in resting-state fMRI studies. Magn. Reson. Med. 68, 1828–1835.

13. Chen, J.E., Polimeni, J.R., Bollmann, S., Glover, G.H., 2019. On the analysis of rapidly sampled fMRI data. Neuroimage 188, 807–820.

14. Chu, P.P.W., Golestani, A.M., Kwinta, J.B., Khatamian, Y.B., Chen, J.J., 2018. Characterizing the modulation of resting-state fMRI metrics by baseline physiology. Neuroimage. https://doi.org/10.1016/j.neuroimage.2018.02.004

15. Cohen, A.D., Chang, C., Wang, Y., 2021. Using multiband multi-echo imaging to improve the robustness and repeatability of co-activation pattern analysis for dynamic functional connectivity. Neuroimage 243, 118555.

16. Cordes, D., Haughton, V.M., Arfanakis, K., Carew, J.D., Turski, P.A., Moritz, C.H., Quigley, M.A., 2001. Frequencies contributing to functional connectivity in the cerebral cortex in “resting-state” data. AJNR Am. J. Neuroradiol. 22, 1326–1333.

17. D Cordes, V M Haughton, K Arfanakis, J D Carew, P A Turski, C H Moritz, M A Quigley, M E Meyerand, 2001. Frequencies contributing to functional connectivity in the cerebral cortex in “resting-state” data. AJNR Am. J. Neuroradiol. 22, 1326–1333.

18. Dipasquale, O., Sethi, A., Laganà, M.M., Baglio, F., Baselli, G., Kundu, P., Harrison, N.A., Cercignani, M., 2017. Comparing resting state fMRI de-noising approaches using multi- and single-echo acquisitions. PLoS One 12, e0173289.

19. Esteban, O., Markiewicz, C.J., Blair, R.W., Moodie, C.A., Isik, A.I., Erramuzpe, A., Kent, J.D., Goncalves, M., DuPre, E., Snyder, M., Oya, H., Ghosh, S.S., Wright, J., Durnez, J., Poldrack, R.A., Gorgolewski, K.J., 2019. fMRIPrep: a robust preprocessing pipeline for functional MRI. Nat. Methods 16, 111– 116.

20. Farras-Permanyer, L., Mancho-Fora, N., Montalà-Flaquer, M., Bartrés-Faz, D., Vaqué-Alcázar, L., Peró-Cebollero, M., Guàrdia-Olmos, J., 2019. Age-related changes in resting-state functional connectivity in older adults. Neural Regeneration Research. https://doi.org/10.4103/1673-5374.255976

21. Fox, M.D., Zhang, D., Snyder, A.Z., Raichle, M.E., 2009. The global signal and observed anticorrelated resting state brain networks. J. Neurophysiol. 101, 3270–3283.

22. Geerligs, L., Renken, R.J., Saliasi, E., Maurits, N.M., Lorist, M.M., 2015. A Brain-Wide Study of Age-Related Changes in Functional Connectivity. Cereb. Cortex 25, 1987–1999.

23. Glasser, M.F., Coalson, T.S., Bijsterbosch, J.D., Harrison, S.J., Harms, M.P., Anticevic, A., Van Essen, D.C., Smith, S.M., 2018. Using temporal ICA to selectively remove global noise while preserving global signal in functional MRI data. Neuroimage 181, 692–717.

24. Glover, G.H., Li, T.Q., Ress, D., 2000. Image-based method for retrospective correction of physiological motion effects in fMRI: RETROICOR. Magn. Reson. Med. 44, 162–167.

25. Golestani, A.M., Chen, J.J., 2022. Performance of Temporal and Spatial Independent Component Analysis in Identifying and Removing Low-Frequency Physiological and Motion Effects in Resting-State fMRI. Front. Neurosci. 16, 867243.

26. Golestani, A.M., Kwinta, J.B., Strother, S.C., Khatamian, Y.B., Chen, J.J., 2016. The association between cerebrovascular reactivity and resting-state fMRI functional connectivity in healthy adults: The influence of basal carbon dioxide. Neuroimage 132, 301–313.

27. Grady, C., Sarraf, S., Saverino, C., Campbell, K., 2016. Age differences in the functional interactions among the default, frontoparietal control, and dorsal attention networks. Neurobiol. Aging 41, 159–172.

28. Griffanti, L., Douaud, G., Bijsterbosch, J., Evangelisti, S., Alfaro-Almagro, F., Glasser, M.F., Duff, E.P., Fitzgibbon, S., Westphal, R., Carone, D., Beckmann, C.F., Smith, S.M., 2017. Hand classification of fMRI ICA noise components. Neuroimage 154, 188–205.

29. Gu, Y., Han, F., Liu, X., 2019. Arousal Contributions to Resting-State fMRI Connectivity and Dynamics. Front. Neurosci. 13, 1190.

30. Hamada, C., Kawagoe, T., Takamura, M., Nagai, A., Yamaguchi, S., Onoda, K., 2021. Altered resting-state functional connectivity of the frontal-striatal circuit in elderly with apathy. PLoS One 16, e0261334.

31. Hausman, H.K., O’Shea, A., Kraft, J.N., Boutzoukas, E.M., Evangelista, N.D., Van Etten, E.J., Bharadwaj, P.K., Smith, S.G., Porges, E., Hishaw, G.A., Wu, S., DeKosky, S., Alexander, G.E., Marsiske, M., Cohen, R., Woods, A.J., 2020. The Role of Resting-State Network Functional Connectivity in Cognitive Aging. Frontiers in Aging Neuroscience. https://doi.org/10.3389/fnagi.2020.00177

32. He, Y., Byrge, L., Kennedy, D.P., 2020. Nonreplication of functional connectivity differences in autism spectrum disorder across multiple sites and denoising strategies. Hum. Brain Mapp. 41, 1334–1350.

33. Hu, S., Chao, H.H.-A., Zhang, S., Ide, J.S., Li, C.-S.R., 2014. Changes in cerebral morphometry and amplitude of low-frequency fluctuations of BOLD signals during healthy aging: correlation with inhibitory control. Brain Struct. Funct. 219, 983–994.

34. Jockwitz, C., Caspers, S., 2021. Resting-state networks in the course of aging-differential insights from studies across the lifespan vs. amongst the old. Pflugers Arch. 473, 793–803.

35. Koch, W., Teipel, S., Mueller, S., Buerger, K., Bokde, A.L.W., Hampel, H., Coates, U., Reiser, M., Meindl, T., 2010. Effects of aging on default mode network activity in resting state fMRI: does the method of analysis matter? Neuroimage 51, 280–287.

36. Li, J., Kong, R., Liégeois, R., Orban, C., Tan, Y., Sun, N., Holmes, A.J., Sabuncu, M.R., Ge, T., Yeo, B.T.T., 2019. Global signal regression strengthens association between resting-state functional connectivity and behavior. Neuroimage 196, 126–141.

37. Liu, T.T., 2016. Noise contributions to the fMRI signal: An overview. NeuroImage. https://doi.org/10.1016/j.neuroimage.2016.09.008

38. Liu, T.T., Nalci, A., Falahpour, M., 2017. The global signal in fMRI: Nuisance or Information? NeuroImage. https://doi.org/10.1016/j.neuroimage.2017.02.036

39. Li, Y.-T., Chang, C.-Y., Hsu, Y.-C., Fuh, J.-L., Kuo, W.-J., Yeh, J.-N.T., Lin, F.-H., 2021. Impact of physiological noise in characterizing the functional MRI default-mode network in Alzheimer’s disease. J. Cereb. Blood Flow Metab. 41, 166–181.

40. Makedonov, I., Black, S.E., Macintosh, B.J., 2013. BOLD fMRI in the white matter as a marker of aging and small vessel disease. PLoS One 8, e67652.

41. Mancho-Fora, N., Montalà-Flaquer, M., Farràs-Permanyer, L., Bartrés-Faz, D., Vaqué-Alcázar, L., Peró-Cebollero, M., Guàrdia-Olmos, J., 2020. Resting-state functional dynamic connectivity and healthy aging: A sliding-window network analysis. Psicothema 32, 337–345.

42. Mazerolle, E.L., Gawryluk, J.R., Dillen, K.N.H., Patterson, S.A., Feindel, K.W., Beyea, S.D., Stevens, M.T.R., Newman, A.J., Schmidt, M.H., D’Arcy, R.C.N., 2013. Sensitivity to white matter FMRI activation increases with field strength. PLoS One 8, e58130.

43. Onoda, K., Yamaguchi, S., 2013. Small-worldness and modularity of the resting-state functional brain network decrease with aging. Neurosci. Lett. 556, 104–108.

44. Parkes, L., Fulcher, B., Yücel, M., Fornito, A., 2018. An evaluation of the efficacy, reliability, and sensitivity of motion correction strategies for resting-state functional MRI. Neuroimage 171, 415– 436.

45. Patil, A.U., Madathil, D., Huang, C.-M., 2021. Healthy Aging Alters the Functional Connectivity of Creative Cognition in the Default Mode Network and Cerebellar Network. Front. Aging Neurosci. 13, 607988.

46. Podgórski, P., Waliszewska-Prosół, M., Zimny, A., Sąsiadek, M., Bladowska, J., 2021. Resting-State Functional Connectivity of the Ageing Female Brain—Differences Between Young and Elderly Female Adults on Multislice Short TR rs-fMRI. Frontiers in Neurology. https://doi.org/10.3389/fneur.2021.645974

47. Power, J.D., Plitt, M., Laumann, T.O., Martin, A., 2017. Sources and implications of whole-brain fMRI signals in humans. Neuroimage 146, 609–625.

48. Pruim, R.H.R., Mennes, M., Buitelaar, J.K., Beckmann, C.F., 2015a. Evaluation of ICA-AROMA and alternative strategies for motion artifact removal in resting state fMRI. Neuroimage 112, 278–287.

49. Pruim, R.H.R., Mennes, M., van Rooij, D., Llera, A., Buitelaar, J.K., Beckmann, C.F., 2015b. ICA-AROMA: A robust ICA-based strategy for removing motion artifacts from fMRI data. Neuroimage 112, 267– 277.

50. Salimi-Khorshidi, G., Douaud, G., Beckmann, C.F., Glasser, M.F., Griffanti, L., Smith, S.M., 2014. Automatic denoising of functional MRI data: Combining independent component analysis and hierarchical fusion of classifiers. NeuroImage. https://doi.org/10.1016/j.neuroimage.2013.11.046

51. Satterthwaite, T.D., Elliott, M.A., Gerraty, R.T., Ruparel, K., Loughead, J., Calkins, M.E., Eickhoff, S.B., Hakonarson, H., Gur, R.C., Gur, R.E., Wolf, D.H., 2013. An improved framework for confound regression and filtering for control of motion artifact in the preprocessing of resting-state functional connectivity data. Neuroimage 64, 240–256.

52. Scheel, N., Tarumi, T., Tomoto, T., Munro Cullum, C., Zhang, R., Zhu, D.C., n.d. Resting-state functional MRI signal fluctuations are correlated with brain amyloid-β deposition. https://doi.org/10.1101/2021.04.22.21255924

53. Schultz, A.P., Chhatwal, J.P., Huijbers, W., Hedden, T., van Dijk, K.R.A., McLaren, D.G., Ward, A.M., Wigman, S., Sperling, R.A., 2014. Template based rotation: a method for functional connectivity analysis with a priori templates. Neuroimage 102 Pt 2, 620–636.

54. Shams, S., LeVan, P., Chen, J.J., 7792764, 2021. The neuronal associations of respiratory-volume variability in the resting state. Neuroimage 230, 117783.

55. Shin, W., Koenig, K.A., Lowe, M.J., 2022. A comprehensive investigation of physiologic noise modeling in resting state fMRI; time shifted cardiac noise in EPI and its removal without external physiologic signal measures. NeuroImage. https://doi.org/10.1016/j.neuroimage.2022.119136

56. Siman-Tov, T., Bosak, N., Sprecher, E., Paz, R., Eran, A., Aharon-Peretz, J., Kahn, I., 2016. Early Age-Related Functional Connectivity Decline in High-Order Cognitive Networks. Front. Aging Neurosci. 8, 330.

57. Song, J., Birn, R.M., Boly, M., Meier, T.B., Nair, V.A., Meyerand, M.E., Prabhakaran, V., 2014. Age-related reorganizational changes in modularity and functional connectivity of human brain networks. Brain Connect. 4, 662–676.

58. Stumme, J., Jockwitz, C., Hoffstaedter, F., Amunts, K., Caspers, S., 2020. Functional network reorganization in older adults: Graph-theoretical analyses of age, cognition and sex. Neuroimage 214, 116756.

59. Tailby, C., Masterton, R.A.J., Huang, J.Y., Jackson, G.D., Abbott, D.F., 2015. Resting state functional connectivity changes induced by prior brain state are not network specific. NeuroImage. https://doi.org/10.1016/j.neuroimage.2014.11.037

60. Thomas, C.G., Harshman, R.A., Menon, R.S., 2002. Noise reduction in BOLD-based fMRI using component analysis. Neuroimage 17, 1521–1537.

61. Tsvetanov, K.A., Henson, R.N.A., Jones, P.S., Mutsaerts, H., Fuhrmann, D., Tyler, L.K., Cam-CAN, Rowe, J.B., 2020. The effects of age on resting-state BOLD signal variability is explained by cardiovascular and cerebrovascular factors. Psychophysiology e13714.

62. Tsvetanov, K.A., Henson, R.N.A., Tyler, L.K., Davis, S.W., Shafto, M.A., Taylor, J.R., Williams, N., Cam-Can, Rowe, J.B., 2015. The effect of ageing on fMRI: Correction for the confounding effects of vascular reactivity evaluated by joint fMRI and MEG in 335 adults. Hum. Brain Mapp. 36, 2248–2269.

63. Van Dijk, K.R.A., Sabuncu, M.R., Buckner, R.L., 2012. The influence of head motion on intrinsic functional connectivity MRI. Neuroimage 59, 431–438.

64. Varangis, E., Habeck, C.G., Razlighi, Q.R., Stern, Y., 2019. The Effect of Aging on Resting State Connectivity of Predefined Networks in the Brain. Front. Aging Neurosci. 11, 234.

65. Whitfield-Gabrieli, S., Nieto-Castanon, A., 2012. *Conn*: A Functional Connectivity Toolbox for Correlated and Anticorrelated Brain Networks. Brain Connectivity. https://doi.org/10.1089/brain.2012.0073

66. Wise, R.G., Ide, K., Poulin, M.J., Tracey, I., 2004. Resting fluctuations in arterial carbon dioxide induce significant low frequency variations in BOLD signal. Neuroimage 21, 1652–1664.

67. Wong, C.W., Olafsson, V., Tal, O., Liu, T.T., 2013. The amplitude of the resting-state fMRI global signal is related to EEG vigilance measures. Neuroimage 83, 983–990.

68. Woo, C.-W., Krishnan, A., Wager, T.D., 2014. Cluster-extent based thresholding in fMRI analyses: pitfalls and recommendations. Neuroimage 91, 412–419.

69. Xie, W., Peng, C.-K., Shen, J., Lin, C.-P., Tsai, S.-J., Wang, S., Chu, Q., Yang, A.C., 2020. Age-related changes in the association of resting-state fMRI signal variability and global functional connectivity in non-demented healthy people. Psychiatry Res. 291, 113257.

70. Yeo, B.T.T., Thomas Yeo, B.T., Krienen, F.M., Sepulcre, J., Sabuncu, M.R., Lashkari, D., Hollinshead, M., Roffman, J.L., Smoller, J.W., Zöllei, L., Polimeni, J.R., Fischl, B., Liu, H., Buckner, R.L., 2011. The organization of the human cerebral cortex estimated by intrinsic functional connectivity. Journal of Neurophysiology. https://doi.org/10.1152/jn.00338.2011

71. Yuan, H., Zotev, V., Phillips, R., Bodurka, J., 2013. Correlated slow fluctuations in respiration, EEG, and BOLD fMRI. Neuroimage 79, 81–93.

72. Zhang, H.-Y., Chen, W.-X., Jiao, Y., Xu, Y., Zhang, X.-R., Wu, J.-T., 2014. Selective vulnerability related to aging in large-scale resting brain networks. PLoS One 9, e108807.

73. Zhong, X.Z., Chen, J.J., 2022. Resting-state functional magnetic resonance imaging signal variations in aging: The role of neural activity. Hum. Brain Mapp. 43, 2880–2897.

